# Epigenomic convergence of genetic and immune risk factors in neurodevelopmental disorder cortex

**DOI:** 10.1101/270827

**Authors:** Vogel Ciernia A., Laufer B.I., Dunaway K.W., Hwang H., Mordaunt C.E., Coulson R.L., Yasui D.H., LaSalle J.M.

**Affiliations:** Department of Medical Microbiology and Immunology, MIND Institute, Genome Center, University of California, Davis, CA 95616

**Keywords:** Neurodevelopmental disorders, Autism spectrum disorders, DNA methylation, microglia, epigenetics

## Abstract

Neurodevelopmental disorders (NDDs) impact 7% to 14% of all children in developed countries and are one of the leading causes of lifelong disability. Epigenetic modifications are poised at the interface between genes and environment and are predicted to reveal insight into the gene networks, cell types, and developmental timing of NDD etiology. Whole-genome bisulfite sequencing was used to examine DNA methylation in 49 human cortex samples from three different NDDs (autism spectrum disorder, Rett syndrome, and Dup15q syndrome) and matched controls. Integration of methylation differences across NDDs with relevant genomic and genetic datasets revealed differentially methylated regions (DMRs) unique to each type of NDD but with shared regulatory functions in neurons and microglia. DMRs were significantly enriched for known NDD genetic risk factors, including both common inherited and rare *de novo* variants. Weighted region co-methylation network analysis revealed a module related to NDD diagnosis and enriched for microglial regulatory regions. Together, these results demonstrate an epigenomic signature of NDDs in human cortex shared with known genetic and immune etiological risk. Epigenomic insights into cell types and gene regulatory regions will aid in defining therapeutic targets and early biomarkers at the interface of genetic and environmental NDD risk factors.

## Introduction

Neurodevelopmental disorders (NDDs) are one of the leading causes of lifelong disability. NDDs including autism spectrum disorders (ASDs), intellectual disabilities, attention-deficit/hyperactivity disorder, cerebral palsy, and Down and fetal alcohol syndrome account for 7% to 14% of all children in developed countries (Miller et al. 2016). The current treatments consist of intensive and expensive behavioral therapy often combined with drug therapies to treat comorbid symptoms such as anxiety. The lifetime cost for a person with ASD, one type of NDD, ranges from $1.2-4.7 million (Buescher et al. 2014) and can cost in caregiver time alone $5.5 million more than for a neurotypical child (Dudley and Emery 2014). Combined with the rising diagnosis rates of ASD and other NDDs and the limited treatments currently available, there is a great need to accelerate the discovery and development of novel NDDs therapeutics and early interventions.

The etiology of NDDs is complex and involves both genetic and environmental risk factors. Rett syndrome (RTT) is an X-linked dominant syndromic ASD/NDD caused by mutations in *MECP2*, affecting females. Duplication 15q11.2-q13.3 (Dup15q) syndrome is a ASD/NDD caused by a maternally inherited copy number variant (CNV) resulting in increased expression of the imprinted gene *UBE3A*. In contrast, cortical idiopathic ASD sample were defined as ASD diagnosis without a known genetic cause. The clinical diversity of idiopathic ASD parallels the genetic complexity of the disorder, which includes hundreds of rare risk variants and potentially thousands of common risk variants (Sanders et al. 2015; De La Torre-Ubieta et al. 2016; The Autism Spectrum Disorders Working Group of The Psychiatric Genomics Consortium 2017). For example, recent exome sequencing approaches in ASD focused on identifying rare, highly penetrant, *de novo* mutations have identified a number of high confidence ASD genes with functions in neuronal synapses and transcriptional regulation (Ben-David and Shifman 2012; Sanders et al. 2015). Genetic and environmental risks are hypothesized to interact in the etiology of ASD (Vogel Ciernia and LaSalle 2016), and a multitude of early-life environmental
perturbations increase the risk of ASD, including maternal immune activation and pollutant exposure during gestation (Lyall, Schmidt, et al. 2014).

Disruption of epigenetic processes regulating brain developmental has been proposed as a potential mechanism linking environmental and genetic risk. For example, transcription, DNA methylation, and histone acetylation analyses in postmortem ASD brain have consistently implicated gene pathways involved in synaptic development and immune function gene pathways (Voineagu et al. 2011; Gupta et al. 2014; Ladd-Acosta et al. 2014; Nardone et al. 2014, 2017; Lin et al. 2016; Parikshak et al. 2016; Sun et al. 2016). However, most analyses of DNA methylation in ASD human brain has been limited to less than two percent of the total CpG sites in the human genome (Ladd-Acosta et al. 2014; Nardone et al. 2014, 2017) or a specific genetic subset of ASD (Dunaway et al. 2016). Consequently, we examined DNA methylation signatures across the entire human genome using unbiased whole-genome bisulfite sequencing (WGBS) of human cortices from three different NDDs and matched controls. Leveraging cortical WGBS data comparing ASD, RTT, and Dup15q with matched controls, we identified regions of differential methylation in loci implicated in genetic risk for NDDs. These loci were enriched in brain developmental functions in both neuronal synapses and microglia. Therefore, differential DNA methylation captures gene pathways and cell types converging with both immune and genetic perturbations in brain development and suggests novel therapeutic pathways for NDDs.

## Methods

### Sample Acquisition, DNA Isolation, and WGBS Library Preparation ASD BA9

Human cerebral cortex samples from Broadmann Area (BA) 9 were obtained from the National Institute of Child Health and Human Development (NICHD) Brain and Tissue Bank for Developmental Disorders at the University of Maryland. DNA was isolated using the QIAGEN Puregene kit (Qiagen, 158667) and WGBS libraries were prepared as described previously (Dunaway et al. 2016). Briefly, 5 μg of DNA was fragmented to ~300 bp using 28 cycles of 15 seconds on/15 seconds off on a Diagenode Bioruptor. DNA was end-repaired using 1× T4 DNA ligase buffer, 400 μM dNTPs, 15 U T4 DNA polymerase (NEB), and 50 U PNK (NEB) for 30 min at 20°C. After PCR purification (Qiagen), adenine bases were appended to the ends using 1× NEB 2 buffer, 200 μM dATP, and 15 U Klenow Fragment (3′ to 5′ exo-, NEB) for 30 min at 37°C. After another DNA purification using the PCR MinElute kit (Qiagen), 3 μL of Illumina’s methylated sequencing adapters were ligated on using 1× ligase buffer and 5 μL Quick T4 DNA Ligase (NEB) for 30 min at room temperature. After a final PCR purification, 500 ng of library was bisulfite converted using Zymo’s EZ DNA Methylation-Direct kit according to the manufacturer’s instructions. The library was then amplified using 2.5 U PfuTurbo Cx Hotstart DNA Polymerase (Stratagene) for 12 cycles using Illumina’s standard amplification protocol. The library’s quality was assessed on a Bioanalyzer (Agilent) and sequenced (100 bp, single-ended) on an Illumina HiSeq 2000. Each biological sample was sequenced on a single lane.

### Sample Acquisition, DNA Isolation, and WGBS Library Preparation Rett BA9

DNA was extracted from BA9 cortex from RTT and Control samples using the Zymo Duet Kit (Zymo). 200ng of total DNA was bisulfite converting using the Zymo Lighting kit (Zymo) and 50ng of converted DNA was used as input for library construction using the Illumina TruSeq DNA Methylation Kit (Illumina). Each sample was given a unique barcode and 14 cycles of PCR amplification. Final libraries were size selected with two rounds of KAPA Pure Beads: 0.7X Left and 0.65X Left/0.55X Right for a final library size centered around 370-470 base pairs. Final libraries were assessed with Bioanalyzer (Agilent), quantified, pooled and run for 150 bp paired end sequencing on two lanes of the HiSeqX (Illumina).

### WGBS Sample Processing

Raw FASTQ files were chastity filtered and then trimmed using trim_galore (https://www.bioinformatics.babraham.ac.uk/projects/trimgalore/) to remove adapters and sequence at the 5′ and 3′ ends with methylation bias. Reads were then aligned to the human genome (hg38), deduplicated, and extracted to a CpG count matrix using Bismark (Krueger and Andrews 2011) QC/QA was performed using Bismark, FastQ screen (Wingett and Andrews 2018), and MultiQC (Ewels et al. 2016). Subsequent analysis was done with custom Perl and R scripts (see github repositories below) (Dunaway et al. 2016).

### Differentially Methylated Regions (DMRs) and Blocks

DMRs were called for diagnosis (NDD versus Control) for each NDD separately using the R package DMRseq (Korthauer et al. 2018). This approach utilizes statistical inference, where smoothed methylation values are obtained from CpG sites that are weighted based on their individual coverage. Bismark cytosine reports were processed to collapse strand-symmetric CpGs using bsseq (Hansen et al. 2012). CpGs from unmapped scaffolds and the mitochondrial sequences were removed from subsequent analysis. For each NDD, the respective samples were filtered to have at least one read of coverage per a CpG across all samples. DMRs were then assembled from correlated CpGs and then a generalized least squares (GLS) regression model with a nested autoregressive correlated error structure for the effect of interest was used to estimate a region statistic for each of the candidate DMRs. DMRs were identified using the default parameters, aside from setting the minimum number of CpGs to 4 per a region and a permutation p-value cutoff of 0.05 for significance testing without additional corrections. For all analyses, the covariate of age was directly adjusted for and for the ASD analysis sex was included as a covariate to balance permutation testing against a pooled null distribution. Background regions for each NDD cohort were defined as all regions that met the criteria for DMR statistical testing. The three background region sets were then merged using bedtools to create a set of consensus background regions, allowing for the use of a consistent background across NDD cohorts. Individual smoothed methylation values for the DMRs were generated using bsseq (Hansen et al. 2012) for visualization. Finally, the blocks of differential methylation were identified using the recommended block parameters from the DMRseq vignette (Korthauer et al. 2018).

### DMR Associated Genes

Genes associated with DMRs or background regions were found using bedtools closest function on all hg38 Ensemble annotated genes and a subsequent filter for genes within +/- 10kb from the start or end of each DMR. With this approach a DMR is first assigned to a gene if it overlaps the gene body. If a DMR does not overlap a gene body then it is assigned based on the closest gene (or genes in case of a tie) upstream or downstream, within the limit of 10kb.

### Principal Components Analysis (PCA)

PCA was performed on average methylation (%mG/CG) over 20kb windows with a minimum coverage of 1 read/CpG and 20 CpGs per window, across the genome using the prcomp and ggbiplot functions in R.

### CIBERSORT Cell Type Deconvolution

Methylation levels from all BA9 cortex samples were extracted for CpGs showing unique cell type specific methylation levels among Glutamaterigc neurons, GABAergic neurons, and glial cells (CpG sites used with at least one read/CpG for all samples, Figures S2) (Kozlenkov et al. 2014). The methylation values from Kozlenkov et al., 2014 were used as custom signature profile for input to CIBERSORT (Newman et al. 2015) (https://cibersort.stanford.edu). CIBERSORT estimated the relative levels of distinct cell types within each WGBS sample by comparing the methylation patterns of the same CpGs between the samples and custom signature profile using machine learning linear support vector regression (Newman et al. 2015).

### Average Methylation Analysis Over Genomic Features

Average methylation was calculated for specific genomic features across all features in the genome for each individual. Genebodies were assigned from transcription start site to transcription end site by the Ensemble hg38 annotation (http://www.ensembl.org/biomart). Promoters were taken from the Ensemble hg38 promoters database (Zerbino et al. 2015) and CpG Islands were taken from the UCSC table browser for hg38 (https://genome.ucsc.edu) (Karolchik et al. 2004). Average methylation levels and average sequencing coverage were analyzed using a mixed model ANOVA with main effects for diagnosis, genomic feature, and sex, the interaction between diagnosis and genomic feature, and the random effect of 1/sampleID. Posthoc comparisons were made with Benjamini-Hochberg corrected t-tests.

### GO Term Enrichment

Gene Ontology (GO) term enrichment on genomic regions was performed for DMRs using the R package GOfuncR (Grote 2018) for GO terms from the current version of the org.Hs.eg.db_3.6.0 package (March, 2018). ASD, Dup15q and RTT DMRs were compared to background consensus regions using a hypergeometric test to compare the overlap between each DMR and gene extended regions (10kb up and downstream of each hg38 Ensemble gene start and end site). The resulting raw over-representation p-values were FDR corrected to p<0.05 and simplified using Revigo (Supek et al. 2011) trimming with default settings for human.

### Region Overlap Enrichment and Datasets

Enrichment of DMRs within specific genomic contexts was performed using Genomic Association Tester (GAT) (Heger et al. 2013) with a workspace defined by the consensus background regions, isochores for hg38 (controls for G+C Bias during random sampling), and 100,000 permutations. Multiple comparisons were Benjamini Hochberg corrected to *p*<0.05. In all figures non-significant enrichments are shown in grey.

Chromatin states were taken from PFC from the ENCODE portal (Ernst et al. 2011; Ernst and Kellis 2013; Roadmap Epigenomics et al. 2015) (http://www.roadmapepiqenomics.org/). DMRs from ASD PFC from 450K array analysis were derived from (Nardone et al. 2017), histone H3K4me3 differential regions in ASD compared to control brains were taken from (Shulha et al. 2012), differential acetylation peaks from ASD brain were derived from (Sun et al. 2016), NeuN+ and NeuN-open chromatin regions in human PFC were taken from (Fullard et al. 2017), Histone marks and PU1 ChIPseq from human microglia were derived from (Gosselin et al. 2017), human PFC NeuN+ and – WGBS DMRs were taken from (Lister et al. 2013), pre versus postnatal DNA methylation data was taken from (Jaffe et al. 2015; Spiers et al. 2015), H3K4me3 across development was taken from (Shulha et al. 2013). Maternal Immune Activation WGBS microglial DMRs were taken from (Vogel Ciernia et al. 2017), and developmental microglial regulatory sites were derived from (Matcovitch-Natan et al. 2016). Human data in hg19 or mouse data was transferred to hg38 using LiftOver (Hinrichs et al. 2006).

### Gene Overlap Enrichment and Datasets

Enrichment for DMR associated genes and previously published gene lists was performed with the GeneOverlaps R package (Shen 2013), where a two-tailed Fisher’s exact test was performed and all overlaps were FDR corrected to *p*<0.05. Gene lists were filtered to include only genes present in the background loci associated genes prior to statistical testing. All gene lists and citations are in Table S7.

#### Briefly, ASD and ID genetic risk genes were taken from SFARI

(https://s1gene.sfari.org/autdb/GSStatistics.do), and recent exome sequencing studies (Gilissen et al. 2014; Iossifov et al. 2014; Sanders et al. 2015). pLI>0.9 genes were identified from the Exome Aggregation Consortium (Lek et al. 2016) as genes with a probability of loss of function mutation > 0.9, indicating that they are highly intolerant to genetic variation in the human population. Human ASD GWAS hits were taken from (The Autism Spectrum Disorders Working Group of The Psychiatric Genomics Consortium 2017) at an association p value < 0.05. Additional GWAS gene lists were taken from https://www.ebi.ac.uk/gwas. Lists of genes were obtained from published datasets for Dup15q and Parikshak et al differentially expressed genes in ASD (Parikshak et al. 2016), Gupta et al differentially expressed ASD genes (Gupta et al. 2014), Gandal et al differentially expressed in ASD brain and other neuropsychiatric disorders (Gandal et al. 2018), RTT differentially expressed genes are from (Lin et al. 2016), genes differentially expressed in Alzheimer’s disease brain (Miller et al. 2013), genes associated with ASD DMRs (Nardone et al. 2017), and genes associated with peaks of differential H3K27 acetylation in ASD brain (Sun et al. 2016). Microglial gene lists were taken from several studies across different microglial isolation approaches and treatments (Hickman et al. 2013; Cronk et al. 2015; Erny et al. 2015; Holtman et al. 2015; Matcovitch-Natan et al. 2016; Hanamsagar et al. 2017; Keren-Shaul et al. 2017; Mattei et al. 2017; Vogel Ciernia et al. 2017; Zhao et al. 2017). Genes with altered expression in offspring following maternal immune activation were also examined from whole brain (Garbett et al. 2012; Oskvig et al. 2012; Richetto et al. 2016).

### Transcription Factor Motif Enrichment

Hypergeometric Optimization of Motif EnRichment (HOMER) (Heinz et al. 2010) was used to identify significantly enriched motifs in NDD DMRs relative to consensus background regions. The findMotifsGenome.pl script was used with corrections for fragment size and CpG content (-chopify –cpg –size given). Available at http://homer.ucsd.edu/homer/

### Microglial Developmental Time course Gene Expression

Raw count data per transcript were downloaded from GEO GSE99622 (Hanamsagar et al. 2017). Two samples were excluded for low total read coverage < 200,000 total reads. The remaining samples were processed in EdgeR to counts per million (CPM) for differential analysis between timepoints and sexes with FDR corrected p-values to 0.05. RPKM values were also calculated by normalizing to gene length for WGCNA.

### Weight Gene Co-Expression Analysis (WGCNA)

WGCNA was performed using the WGCNA R package (version 1.61 2017-08-05) (Langfelder and Horvath 2008, 2012). Reads per kilobase per million (RPKM) data were for dorsal lateral prefrontal cortex samples were taken from the human BrainSpan dataset (http://www.brainspan.org/rnaseq/) (Miller et al. 2014). FPKM values were calculated as described above for microglial development data with the exclusion of LPS treated samples (Hanamsagar et al. 2017), and mouse ensemble ids were converted into human using BioMart. Genes with zero variance in expression or a median absolute deviation of zero were removed from the analysis. Genes with a minimum RPKM of 0.25 or higher in at least one sample were kept for analysis for a total of 25,129 genes (unique Ensemble IDs) for BrainSpan and 12,597 for the microglia dataset (Hanamsagar et al. 2017). Values were transformed to log_2_(RPKM+1) and clustered to visualize outliers. There were no outlier samples in the BrainSpan data but two samples were removed from the microglial development dataset due to poor clustering (Figure S9). A correlation matrix using biweight midcorrelation between all genes was then computed for all samples. An estimated soft thresholding power was used to derive a signed adjacency matrix with approximately scale-free topology (R^2 fit indices >.80) that was then transformed into a topological overlap matrix (TOM). The matrix 1-TOM was used as input to calculate co-expression modules with hierarchical clustering and a minimum module size of 200 genes. The resulting module eigengenes were clustered based on their correlation and modules were merged at a cutheight of 0.25 (correlation of .75) to produce co-expression modules and one additional model with genes that did not show module membership (grey). Pearson’s correlation coefficients were used to calculate the correlation between sample traits and module membership. The expression profile of each module was further summarized by the module eigengene (ME), the first principle component of the module. Intramodule connectivity (kME) was calculated as the correlation between every gene in the module with the module ME. Overlap between module genes and gene lists for NDD genetic risk, NDD DMR associated genes, and NDD differentially expressed genes was calculated using the enrichmentAnalysis function in the anRichment R package (version 0.82-1). The background was set to the intersection between genes in the analysis and the organism database (org.Hs.eg.db_3.4.0) and a Fisher’s Exact test was conducted for each module-gene list pair and corrected to an FDR of p = 0.05. Cell type enrichment was performed similarly with custom lists of cell type gene signatures identified in human brain single cell RNAseq from multiple datasets (Habib et al. 2017; Lake et al. 2018; Zhong et al. 2018). GO term enrichment was similarly calculated using enrichmentAnalysis with the current hg38 human GO term database.

### Weight Region Co-Methylation Network Analysis (WRCMNA)

WRCMNA was performed using the WGCNA R package (version 1.61 2017-08-05) (Langfelder and Horvath 2008, 2012) on smoothed mCpG values extracted for all background consensus regions identified in the DMR analysis across all NDDs. Regions were filtered for a least 1x coverage and the resulting 92836 regions were used as input to the network. A biweight midcorrelation matrix was constructed for all regions and estimated soft thresholding power of 14 was used to derive a signed adjacency matrix with scale-free topology (R^2 fit indices > .8). This matrix was then transformed into a topological overlap matrix (TOM), and the matrix 1-TOM was used as input to for blockwise construction of the co-methylation modules with hierarchical clustering and a minimum module size of 200. The methylation profile of each module was further summarized by the module eigengene (ME), the first principle component of the module. The MEs from the resulting 13 modules and the additional grey module (nonclustered regions) were then correlated to NDD diagnosis, sex, sequencing run, brain region, PMI and age. The resulting Pearson’s correlation p-values were FDR corrected to p=0.05.

Intramodule connectivity (kME) was used to identify top “hub” regions within each module. kME was calculated as the correlation between the absolute values of region significance (the association of individual regions with diagnosis) and module membership (correlation between module eigengenes and smoothed methylation levels). Two-tailed Fisher’s exact tests with FDR correction were used to assess the enrichment of each set of module regions with regulatory regions identified from ENCODE chromHMM for PFC, regions of differential chromatin accessibility and methylation within NeuN+ neuronal compared to NeuN-non-neuronal (microglia, astrocytes, etc.) (Lister et al. 2013; Fullard et al. 2017), and regulatory regions identified from acutely isolated human brain microglia (Gosselin et al. 2017). The background was set as all regions included in the complete network. GO term enrichment was performed on module regions using GOfuncR as described above with background regions set to the complete network.

## Results

### Differentially methylated regions identified in NDD cortex

To identify an epigenomic signature for NDDs, we examined genome-wide methylation profiles from postmortem cortices from three different NDDs and matched controls. WGBS was performed on a total of 49 cortical samples (both previously published and newly generated), and then analyzed through a standardized bioinformatic pipeline and quality control assessment (Table S1 and Figure S1). We compared WGBS methylation levels between brain samples from cortical region BA9 from donors diagnosed with idiopathic ASD (n=12 M and 5 F ASD versus n=5 M and 5 F control), RTT (n=6 F RTT versus n=6 F control), and BA19 from Dup15q syndrome (n=5 M versus n=5 M control) (Dunaway et al. 2016) (Table S1). Global levels of percent methylation at CpG sites (mCpG) and CpH sites (mCpH) trended towards lower levels globally in the ASD compared to control samples, but did not reach statistical significance (p<0.07). In contrast, no global differences in methylation were observed for the RTT samples compared to controls (Figure S1, Table S1). Consistent with previously published findings (Dunaway et al. 2016), hypomethylated global levels of mCpG were observed in Dup15q (Dup15q < Ctrl, p=0.05). No significant differences in mCpG levels were observed over several types of genomic features including CpG islands, gene bodies, or promoters between any of the NDDs and matched controls (Figure S2). Cell type deconvolution with CIBERSORT (Newman et al. 2015) using cell type specific methylation data from sorted human glutamatergic neurons, GABAergic neurons, and glial cells (Kozlenkov et al. 2014) followed by repeated measures ANOVA did not reveal any significant gross cell type differences among any of the NDD cohorts (Figure S3, Table S2). Average mCpG levels within 20 kb windows revealed strong batch effects between NDD sequencing cohorts (disparate brain regions and sequencing platforms), but not between sexes or diagnosis within cohorts (Figure S3). A large domain (69 Mb) of hypomethylation (Dup15q<Control) previously identified in chromosome 15 in Dup15q brain WGBS (Dunaway et al. 2016) and an additional novel region of hypomethylation (23 Mb) on chromosome 16 (Figure S3), indicative of the large scale change in genome CpG methylation in Dup15q were observed. No large megabase block regions were identified in either the ASD or RTT samples.

We next identified differentially methylated regions (DMRs) for each of the three NDDss compared to age matched controls. This was done using the DMRseq R package (Korthauer et al. 2018) with direct adjustment for the covariate of age (all cohorts) and a balancing of permutations for sex (ASD cohort). DMRseq identified 378 significant (permutation *p* < 0.05) DMRs with lower methylation (hypomethylated) and 142 regions with higher methylation (hypermethylated) in ASD compared to control cortices (Figure 1A and Table S3). A large proportion of the ASD DMRs were found on chromosome X (123/520, 24%). While these X chromosome DMRs show a different overall level of methylation between sexes, the relative difference between ASD and matched controls remains significant across the sexes, indicating that sex is an important variable for consideration in the disorder and highlights the importance of including sex as a biological covariate for balanced permutations in our DMR analysis. For the Dup15q cohort, 1750 hypermethylated DMRs (Dup15q>Ctrl) and 1129 hypomethylated DMRs (Dupr15q<Ctrl) were identified as significant (permutation *p* < 0.05) (Figure 1B). Comparing the RTT cohort to controls identified 2730 hypermethylated (RTT>Ctrl) and 1911 hypomethylated (Ctrl>RTT) significant DMRs (Figure 1C) (permutation *p* < 0.05). The locations of the NDD DMRs were largely non-overlapping among the three different conditions (Figure S4 and Table S3).

**Figure 1.**
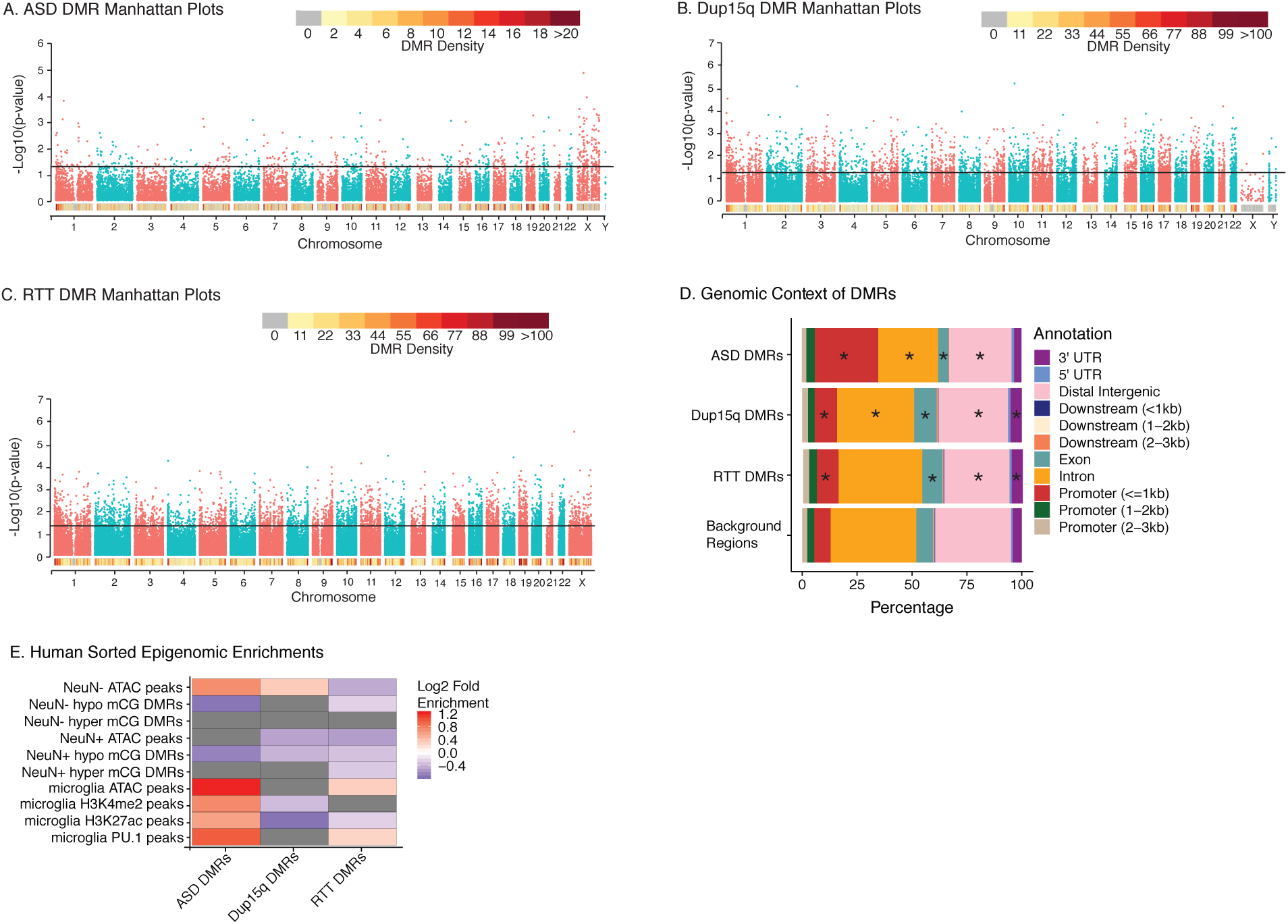
Locations and characteristics of regions of differentially methylated regions (DMR) in ASD, Dup15q, and RTT cortex. **A.** Manhattan plot for DMRs by chromosome for ASD versus controls. Black line represents significance cutoff level for permutation p-values. **B.** Manhattan plot for DMRs by chromosome for Dup15q versus controls. Black line represents significance cutoff level for permutation p-values. **C.** Manhattan plot for DMRs by chromosome for RTT versus controls. Black line represents significance cutoff level for permutation p-values. **D.** DMR enrichment within genomic features relative to background regions. UTR, untranslated region. ^∗^p< 0.05 FDR corrected two-tailed Fisher’s exact test. Table S3 includes statistics. **E.** Enrichment for DMRs over epigenomic profiles from sorted human brain cell populations. ASD DMRs were significantly enriched for regulatory regions active in neurons (NeuN+) and nonneuronal cells (NeuN-), specifically microglial regulatory regions (Lister et al. 2013; Fullard et al. 2017; Gosselin et al. 2017). Color scale reflects log_2_ fold enrichment relative to background regions (p<0.05, GAT permutation test with Benjamini Hochberg correction), and grey represents non-significant differences. Statistic details are shown in Table S5.

ASD, Dup15q, and RTT DMRs were all significantly (Fisher’s exact test FDR p<0.05) enriched within promoter regions near the TSS and de-enriched in intergenic regions compared to background regions (defined in Methods). Dup15q and ASD DMRs were also both de-enriched within introns. RTT and Dup15q DMRs were positively enriched while ASD DMRs were de-enriched for exons (Figure 1D and Table S3). Gene Ontology (GO) enrichment analysis based on genomic locations for ASD DMRs did not reveal any significant GO terms that passed FDR correction. However, top ranked, uncorrected p-values indicate a number of different biological process may be impacted in ASD including vocalization behavior, embryonic patterning, and nervous system development (Figure S5, Table S4). Enrichment for Dup15q DMRs (FDR p<0.05) revealed significant GO terms for processes related to nervous system development including calcium ion binding, cytoskeletal protein binding, neurogenesis, and cell projection, similar to those previously described (Dunaway et al. 2016). RTT DMRs were additionally enriched (FDR p<0.05) for GO terms for calcium ion binding and cytoskeletal protein binding, as well as terms involved in cellular adhesion, system development, plasma membrane, and synapses. Together, these analyses suggest that general nervous system development, and neuronal function in particular, is reflected in brain methylation differences in NDDs.

To further evaluate cell type specificity and regulatory functions, DMRs were overlapped with chromatin state maps from human prefrontal cortex (PFC) (Ernst et al. 2011; Ernst and Kellis 2013; Roadmap Epigenomics et al. 2015), revealing significant (hypergeometric permutation testing FDR p<0.05) enrichment for promoter regions for all three NDD DMRs, and bivalent enhancers for Dup15q and RTT DMRs (Figure S4). ASD, Dup15q and RTT DMRs were de-enriched for regions of heterochromatin, weak transcription, quiescence and several subtypes of enhancers (Figure S6A and Table S5). DMRs were next examined for enrichment for differential regions of chromatin accessibility and methylation within specific cell populations, including neurons (NeuN+) compared to non-neurons (NeuN-, microglia, astrocytes, etc.) isolated from postmortem PFC (Lister et al. 2013; Fullard et al. 2017) and regulatory regions identified from acutely isolated human brain microglia (Gosselin et al. 2017) (Table S5). ASD DMRs were significantly enriched (FDR p<0.05) for regions with marks of open chromatin (ATAC peaks) in non-neuronal cells (Figure 1E), while Dup15q DMRs showed a less significant enrichment for non-neuronal ATAC peaks, while RTT DMRs were significantly de-enriched for similar regions. ASD DMRs were also positively enriched for multiple microglial regulatory elements (Gosselin et al. 2017) including ATAC peaks, active transcription or enhancer regions (H3K4me2 and H3K27ac) and binding sites for the microglial lineage determining transcription factor PU.1 (FDR p<0.05). RTT DMRs showed significant enrichment for microglial ATAC peaks and PU.1 binding. In contrast, Dup15q DMRs were significantly de-enriched for both microglial histone marks (Figure 1E) (FDR p<0.05).

At least some of the regulatory elements found within DMRs may be impacted early in human brain development, as both RTT and Dup15q DMRs showed significant enrichment (FDR p<0.05) for DMRs responsive to changes in early brain development (Spiers et al. 2015) (Figure S6). A direct comparison of the DMRs to previously identified array based methylation studies in ASD brain did not show any significant overlap (Nardone et al. 2017) (Table S5). ASD DMRs, but not Dup15q or RTT DMRs, were enriched for regions identified from prior studies as exhibiting decreased H3K4me3 levels (Shulha et al. 2012) in ASD brain (Figure S6) (FDR p<0.05).

### ASD DMR associated genes are enriched for common and rare NDD genetic risks

To examine overlap between genetic risk factors and epigenetic signatures for NDDs, DMRs were assigned to the closest gene (within the gene body or within +/-10kb of the gene start or end sites, see Methods). ASD DMRs were associated with 478 genes, Dup15q DMRs with 2286 genes and RTT DMRs with 3464 genes (Table S6). Genes associated with each set of NDD DMRs significantly overlapped (two-tailed fisher’s exact test FDR p<0.05) between all three NDDs examined (Table S7 and Figure 2A) and both RTT and Dup15q DMR genes were significantly enriched for genes associated with ASD DMRs identified by a previous 450k array analysis (Nardone et al. 2017) (Table S7). To assess the potential convergence of differential methylation with genetic risk, we overlapped the DMR associated genes from each of the three NDDs with established genetic risk factors for ASD and intellectual disability (Sanders et al. 2015) (Figure 2B, Table S7). ASD, Dup15q, and RTT DMR associated genes were significantly enriched (FDR p<0.05) for multiple curated lists of known risk genes for both ASD and ID.

**Figure 2.**
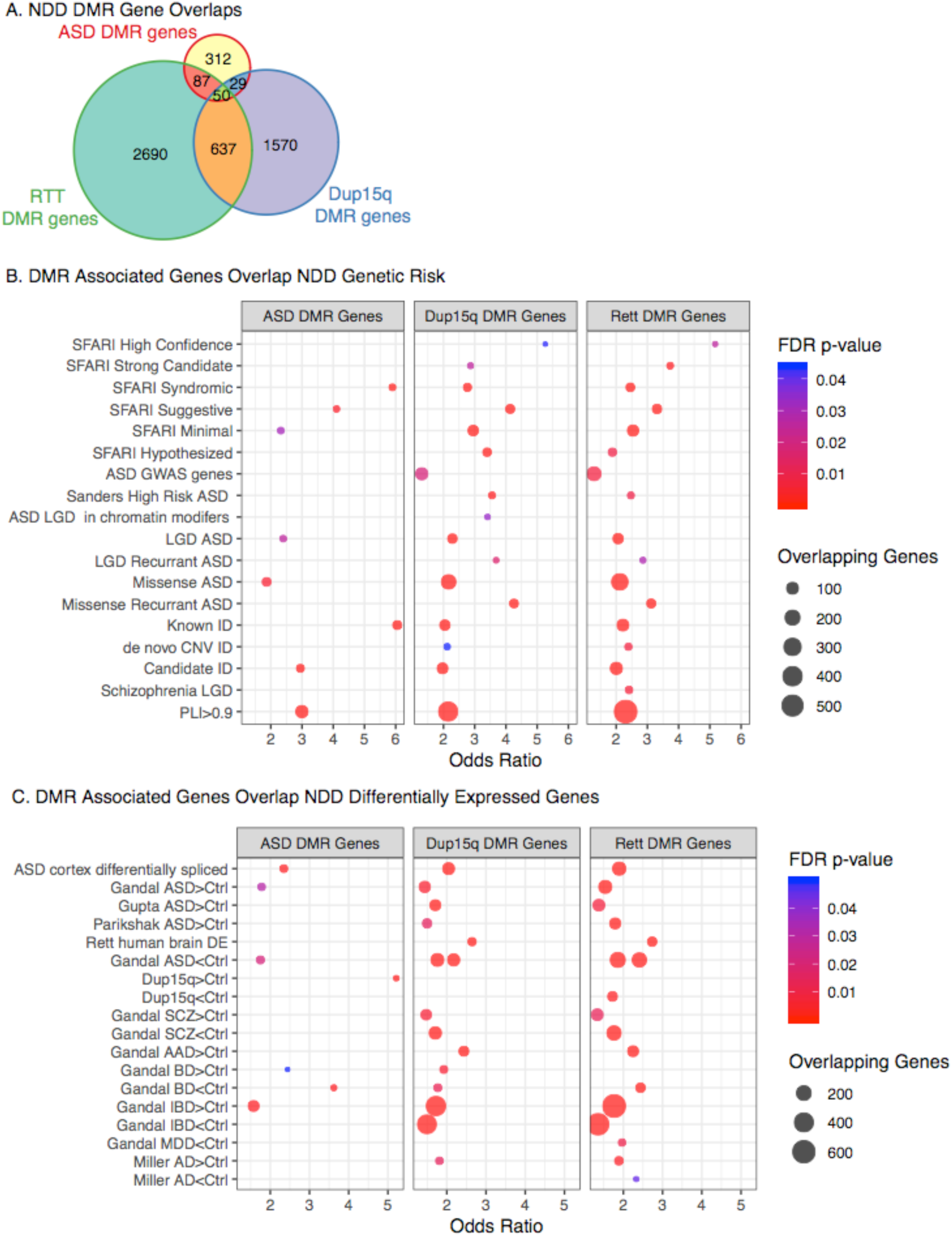
Enrichment of DMR associated genes for NDD genetic risk and differentially expressed genes. **A**. ASD, Dup15q and RTT DMR associated genes significantly overlap with each other (all overlaps are FDR corrected p< 0.05, statistics in Table S7). **B.** ASD, Dup15q, and RTT DMR associated genes are significantly enriched for genes associated with elevated genetic risk for ASD, intellectual disability (ID) or schizophrenia. **C.** ASD, Dup15q, and RTT DMR associated genes are significantly enriched for genes with altered gene expression in numerous neurodevelopmental, neuropsychiatric and neurodegenerative disorders. Significant overlaps are shown from FDR corrected p<0.05 two tailed Fisher’s exact test. SFARI: Simons Foundation Autism Research Initiative, LGD: likely gene disrupting mutation, ID: intellectual disability, SCZ: Schizophrenia, AD: Alzheimer’s Disease, BD: Bipolar Disorder, IBD: Irritable Bowel Syndrome, AAD: Alcohol Abuse Disorder, MDD: Major Depressive Disorder, pLI>0.9: probability of being loss-of-function >0.9 (top genes that are intolerant to human mutation), DE: differentially expressed, Rett human brain DE: DE genes from (Lin et al. 2016), Dup15q: Chromosome 15q11.2-q13.1 duplication syndrome. Gandal: DE genes from (Gandal et al. 2018), Gupta: DE genes from (Gupta et al. 2014), Parikshak: DE genes from (Parikshak et al. 2016), Miller: DE genes from (Miller et al. 2013). Table S7 contains all gene lists, citations and statistics.

These overlaps included genes for multiple types of mutations associated with ASD risk including probable gene disrupting mutations, missense mutations (Iossifov et al. 2014), and genes curated by SFARI. Gene lists associated with other non-NDD traits and disorders from the National Human Genome Research Institute and European Bioinformatics Institute GWAS Catalog (MacArthur et al. 2017) were used as a specificity comparison (Table S6). The vast majority of GWAS targets for other disorders did not show significant enrichment (2/294) with the only exceptions being enrichment for RTT DMR associated genes with genes identified for height or systemic lupus (Table S7).

All three groups of NDD DMR associated genes were enriched for genes identified as subject to strong selection against mutations, indicating that NDD DMRs are impacting genes critical for brain function. To further relate NDD DMRs to genes related to brain function, we overlapped DMR associated genes with genes identified as differentially expressed in neurodevelopment (ASD), neuropsychiatric (schizophrenia, alcoholism, major depressive disorder and bipolar disorder) and neurodegenerative (Alzheimer’s disease) disorders. All three NDD DMR types were significantly enriched (two-tailed Fisher’s exact test FDR p<0.05) for genes with altered expression in brain from ASD, schizophrenia, and bipolar disorder (Figure 2C and Table S7). At least a subset of these overlapping genes may be driven by neuroinflammatory signatures, as there was also a significant overlap between all three NDD DMR associated gene sets and genes with altered expression from patients with irritable bowel syndrome, a disorder characterized by chronic inflammation. Interestingly, RTT and ASD DMR associated, but not Dup15q DMR associated, genes significantly overlapped transcripts showing altered expression in brain from Dup15q. RTT and Dup15q DMR genes, but not ASD DMR genes, were additionally enriched for genes with altered expression in Alzheimer’s disease, RTT brain, and several ASD differential expression lists (Figure 2C and Table S7) (FDR p<0.05). All three NDD DMR associated gene lists were enriched for genes with differential splicing in ASD brain (Parikshak et al. 2016) and genes involved in chromatin regulation, including several splicing factors, suggesting that NDD DMRs may be related to alterations in transcriptional regulation. The differences in overlapping gene lists may reflect differences in the underlying etiology of the genetic NDDs compared to the more heterogenous ASD samples.

### NDD DMR associated genes are dynamically regulated across human brain development

To further examine when NDD DMR genes are most transcriptionally active during brain development, we utilized weighted gene co-expression networks (WGCNA) (Langfelder and Horvath 2008) built from prefrontal cortex brain developmental stages using the human BrainSpan atlas (Miller et al. 2014) (Figure S5 and Table S8). Twelve modules were identified (Figure S7), four were significantly correlated (FDR p value <0.05), and two marginally correlated (FDR < 0.08) with developmental age (Figure 3 and Figure S7). ASD DMR associated genes were not significantly enriched within any of the modules, but were most often found within the midnight blue module (uncorrected p<0.07) (Table S8). Dup15q DMR genes were significantly enriched within three modules (blue, midnight blue and pink) and RTT DMR genes were significantly enriched within blue, magenta, midnight blue, pink and purple modules (FDR p<0.05) (Figure 3B). Of the modules enriched with NDD DMR associated genes, all but the purple module showed dynamic developmental regulation across brain development, indicating that genes impacted by NDD DMRs are critically regulated during brain development. In addition, several of the modules enriched for NDD DMR genes are also enriched for genes differentially expressed in brain disorders or associated with certain types of NDD genetic risk factors. For example, the blue module, which contains genes associated with both Dup15q and RTT DMRs, was also enriched for genes with increased expression in Dup15q, ASD, schizophrenia, bipolar disorder, and Alzheimer’s disease brain samples. In comparison, the midnight blue module was enriched for RTT and Dup15q DMR genes (FDR p<0.05) and marginally for ASD DMR genes (uncorrected p<0.07). This module shows a pattern of increasing expression during mid and late pregnancy and enrichment for genes associated with decreased expression in ASD and schizophrenia, as well as several genetic risk factors for ASD, suggesting that NDD DMRs and genetic risk factors may critically impact third trimester brain development. Modules enriched for NDD DMR associated genes were also enriched for a number of cell type specific markers and processes indicating that multiple types of neuronal and non-neuronal cell types are likely impacted across brain development. For example, the midnight blue module was significantly enriched for multiple types of neuronal and non-neuronal cells, indicating that disruption of genes within this module may alter developmental trajectories across a number of important brain processes.

**Figure 3.**
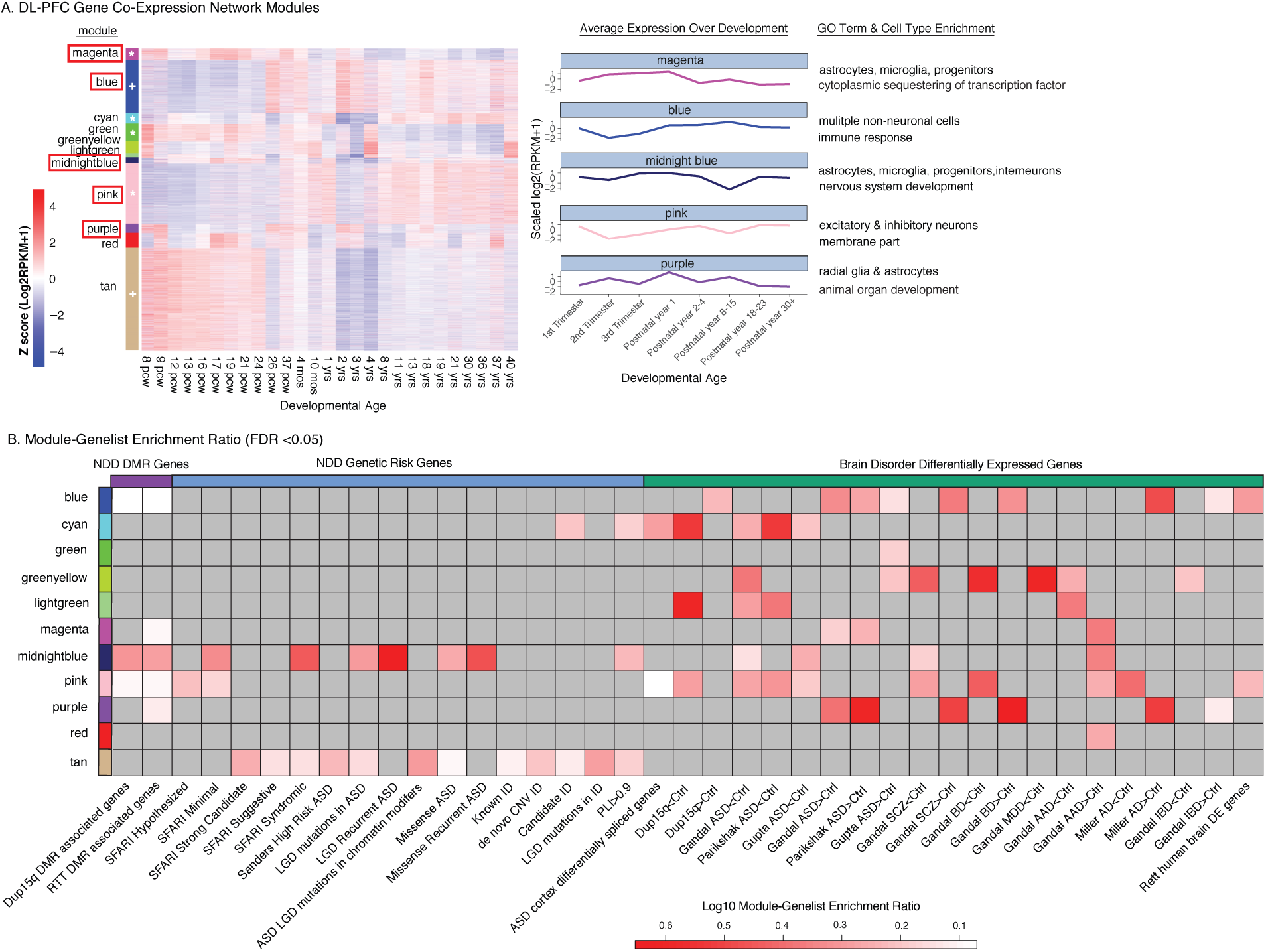
NDD DMR associated genes are enriched in developmentally regulated gene coexpression modules. **A.** Gene expression within each module identified by WGCNA across human DL-PFC brain development. Data from BrainSpan (Miller et al. 2014). Modules with module epigengenes (ME) correlated with developmental age are denoted with **^∗^** for FDR p<0.05 or **+** for FDR p<0.08 (left side color bar). Modules with significant enrichments for any of the three NDD (ASD, Dup15q, or RTT) DMR associated genes are boxed and shown as average gene expression across development time (right panel). Modules relevant to NDD DMRs are labeled with cell type enrichment and/or their top enriched GO term (all enrichments are shown in Table S8). pcw: post-conception week. **B.** Fisher’s exact tests for gene list enrichment within each co-expression module was performed, with results summarized as a heat map of Log10 enrichment (grey, not significant), with all statistical results shown in Table S8. Gene lists are further described with citations in Table S8.

### NDD DMRs overlap microglial genes critical for development and environmental responses

In order to better understand phenotypes associated with the immune signature observed in Figure 1G and previously in ASD brain gene expression datasets (Voineagu et al. 2011; Gupta et al. 2014; Gandal et al. 2016; Lin et al. 2016; Parikshak et al. 2016), we explored the potential overlap of NDD DMR associated genes with several mouse microglial expression datasets. These included microglial developmentally regulated genes (Matcovitch-Natan et al. 2016; Hanamsagar et al. 2017) as well as genes misregulated in microglia isolated from both genetic and maternal immune challenge ASD models (*Mecp2* mutation, PolyI:C maternal immune activation, maternal allergic asthma). We also compared NDD DMR genes to those gene transcripts altered in mouse microglia from germ-free conditions, immune stimulation (LPS, primed, sensome), or an Alzheimer’s model with known disease associated microglia (DAM) involvement (Table S9). All three NDD DMR associated gene lists showed significant enrichment (two-tailed Fisher’s exact test FDR p<0.05) for genes regulated during microglial development, especially genes differentially regulated between later embryonic and postnatal timepoints both in male and female microglia (Figure 4A). NDD DMR associated genes were also enriched for genes misregulated in microglia in several NDD mouse models including genes with altered expression in microglia from a maternal allergic asthma model (MAA), PolyI:C maternal immune activation (MIA) model, and the MeCP2 heterozygous and null mouse models of RTT. The RTT and Dup15q DMR genes significantly overlapped DMRs identified in adult prefrontal cortex for animals with MIA on gestational day (GD) 17 or 9, but the ASD DMR genes only overlapped with the GD17 treatment condition, indicating that the genes impacted by ASD DMRs may be more sensitive to alterations in immune function later in pregnancy. All three NDD DMR sets also overlapped genes normalized by minocycline treatment in adult mice that had MIA induced *in utero* (Figure 4B), suggesting possible benefit of minocycline treatment in human NDDs. NDD DMR associated genes were also enriched for several markers of microglial inflammation and dysfunction during Alzheimer’s disease progression (Figure 4B).

**Figure 4.**
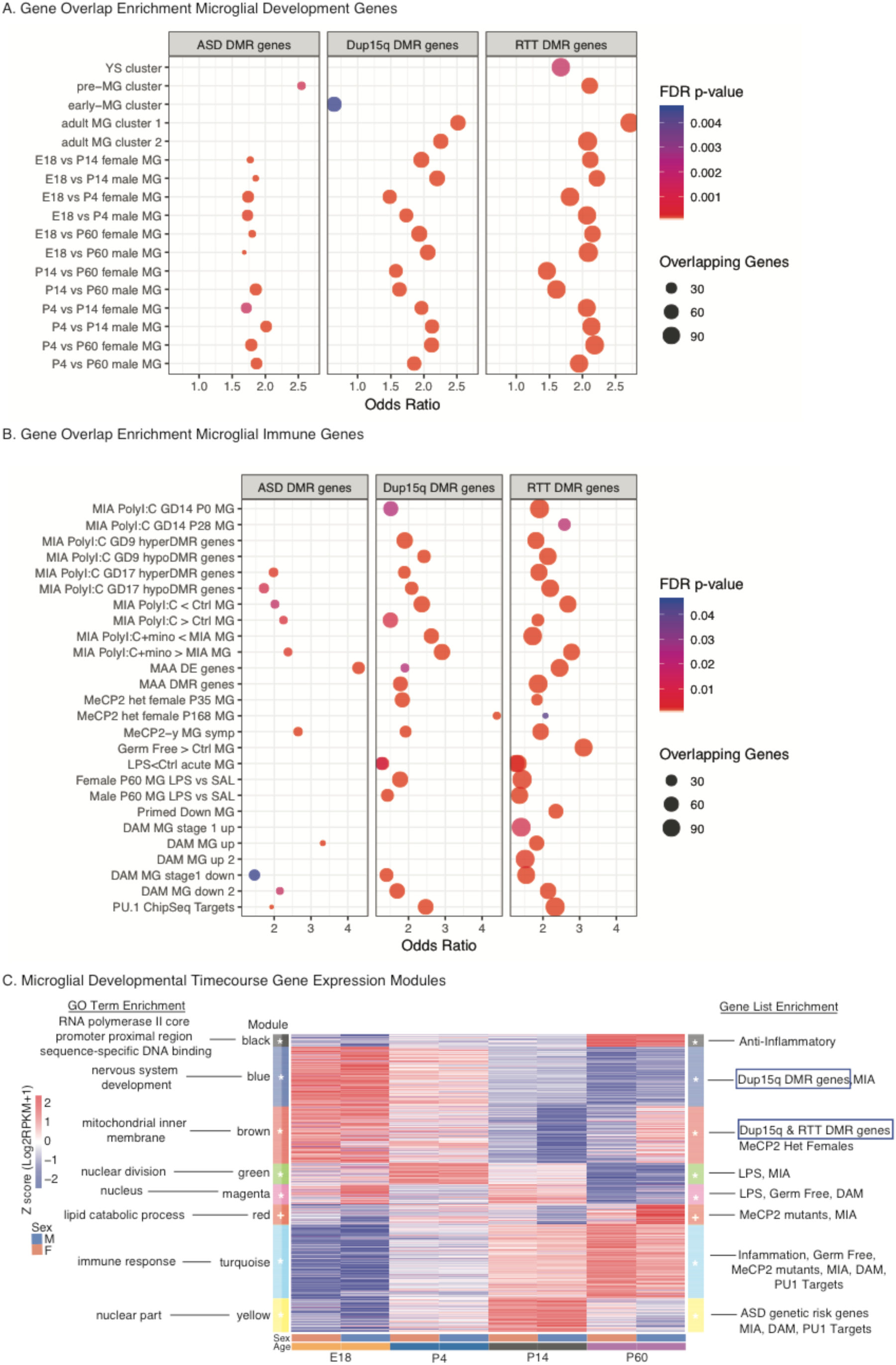
NDD DMR associated genes are enriched for functions in microglial development and regulation. **A.** NDD DMR associated genes were significantly enriched for genes showing developmental regulation during microglial (MG) development from two independent mouse RNA-seq datasets (Matcovitch-Natan et al. 2016; Hanamsagar et al. 2017). All mouse genes were converted to hg38 using BioMart. **B.** NDD DMR associated genes were also significantly enriched for genes regulated in microglia in response to maternal immune activation (Matcovitch-Natan et al. 2016; Mattei et al. 2017; Vogel Ciernia et al. 2017), MeCP2 mutation in heterozygous (MeCP2 het female (Zhao et al. 2017)), or hemizygous (MeCP2-/y (Cronk et al. 2015)) mice, germ free conditions (Erny et al. 2015), LPS versus saline injection (LPS vs SAL) (Holtman et al. 2015; Hanamsagar et al. 2017), primed microglia (Holtman et al. 2015), Alzheimer’s disease associated microglia (DAM) (Keren-Shaul et al. 2017), or the characterized targets of the microglial lineage determining transcription factor PU.1 (Satoh et al. 2014). ^∗^FDR corrected p<0.05 two tailed Fisher’s exact test. Statistical detail and citations are shown in Table S9. **C.** Weighted Gene Co-Expression Network analysis (WGCNA) of microglial development in male and female mice (GEO GSE99622) (Hanamsagar et al. 2017). Normalized gene expression is shown for each identified module in WGCNA in Figure S6. The top enriched gene ontology term for each module is shown on the left. Enriched microglial gene lists are shown on the right (additional data in Table S10 and Figure S6). Modules with significant associations with NDD DMR genes are boxed in blue. ^∗^p<0.05 and +p<0.08 indicate module eigengenes correlated with developmental time (Table S10) (left side color bar). E18: embryonic day 18, P4: postnatal day 4, P14: postnatal day 14, P60: postnatal day 60 (adult). Gene lists, statistics and citations are provided in Table S10.

To identify specifically when NDD DMR genes are most transcriptionally active in microglia, we performed WGCNA on a published RNAseq dataset (Hanamsagar et al. 2017) from microglia isolated from four developmental timepoints from both male and female mice (Figure 4C, Figure S8, and Table S10). Microglia co-expression modules were enriched for GO terms related to microglial function and development as well as differentially expressed genes from microglia studies examining inflammation, genetic mutations and microglial development (Figure S7 and Table S10). ASD genetic risk variants were predominantly enriched (Fisher’s exact test FDR p<0.05) within the yellow module of postnatal day 14 microglial expressed genes. Yellow module microglial genes were predominantly nuclear, PU.1 targets, and misregulated in microglia from both MIA and DAM mouse models. In contrast, Dup15q DMR associated genes were significantly enriched (FDR p<0.05) for the blue module, which was characterized by embryonic and early postnatal microglial expression, nervous system development, dysregulation in MIA microglia, and decreased expression in several human brain disorders including ASD (Table S10). Both Dup15q and RTT DMR genes were significantly enriched (FDR p<0.05) within the brown module, which was also expressed most highly in early microglial development and was enriched for mitochondrial function and energy regulation (Figure 4C and Table S10). The ASD DMRs were not significantly enriched for any individual module (all FDR p>0.05), but approached significance for the brown module (FDR p=0.13). Together, these results indicate a genetic NDD risk within postnatally expressed microglial genes plus an epigenetic NDD signature over genes involved in microglia-neuron interactions and metabolism early in microglial development.

### NDD DMRs are enriched for transcription factor binding sites critical for neurons and microglia

To further evaluate the potential impact of altered DNA methylation on gene regulation, we examined NDD DMRs for known transcription factor binding motif enrichment compared to background regions using HOMER (Heinz et al. 2010). RTT DMRs were enriched for 33 unique transcription factor motifs, 27 of which passed FDR correction (FDR p<0.05). ASD DMRs were enriched for 4 unique motifs and Dup15q DMRs for 7 motifs, none of which passed FDR correction (Figure 5A and Table S11). To assess which cell types might be most impacted by altered transcription factor binding, expression of transcription factor genes was compared across cell types (Lake et al. 2018) and prenatal PFC development (Zhong et al. 2018) from human single cell RNAseq (Figure 5B and Figure S9), with a subset revealing preferential cell type specific expression patterns. For example, the motif for *MEF2D* was enriched in both Dup15q and RTT DMRs and was highly expressed in several excitatory and inhibitory neuronal subtypes and in microglia during early prenatal development. A number of transcription factors impacted by RTT DMRs were most highly expressed at the end of gestation in GABAergic neurons (*HOXA10, HNF1B, PIT1, PAX7*, and *CDX2*), indicating this may be a critical timepoint for cell type specific impacts of RTT DMRs on transcription factor binding and gene regulation. Both Dup15q and RTT DMR-enriched transcription factors *SPI1* (PU.1) and *IRF8* showed preferentially high expression in microglia and developmental regulation in human prenatal PFC. Together, these results demonstrate the potential relevance of changes in DNA methylation for transcription factor binding within several neuronal and non-neuronal cell types across developmental disorders.

**Figure 5.**
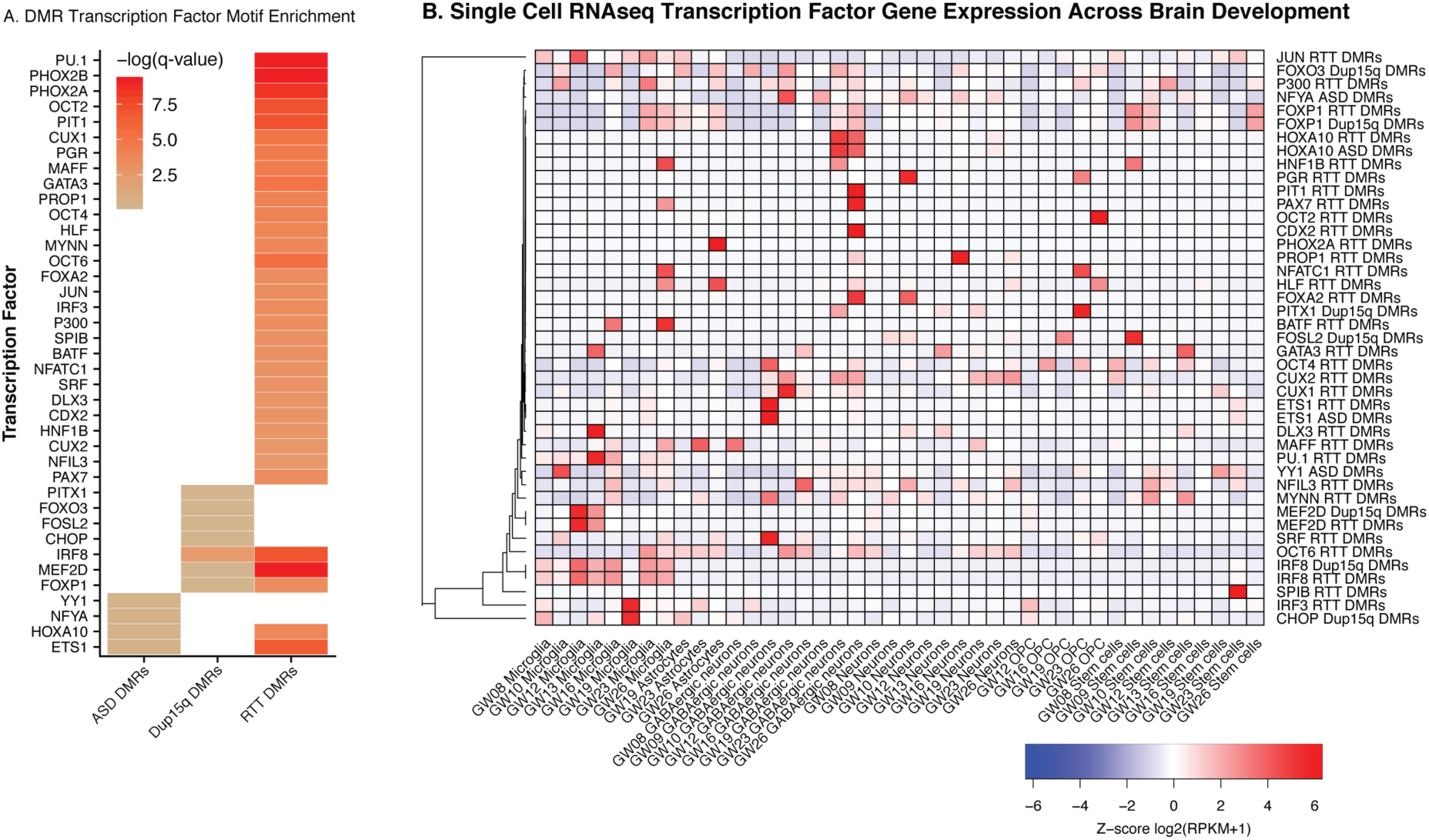
NDD DMRs are enriched for binding motifs for transcription factors regulated during human brain development. **A.** Transcription factors with significantly enriched motifs in NDD DMRs are shown as colors by FDR adjusted log q values (white represents nonsignificant). Enrichments are included for all HOMER binding motifs with uncorrected p<0.05. **B.** Expression of enriched transcription factors (from A) across cell types in human prenatal prefrontal cortex development (single cell RNAseq data from (Zhong et al. 2018)). Expression values shown as Z-scores (within row normalization) of log_2_(RPKM +1) values for each cell type across gestational weeks (GW). Each transcription factor gene is also labeled with the type of enriched NDD DMR (ASD, Dup15q, or RTT). Table S11 includes results of statistical tests. GW: gestational week, OPC: oligodendrocyte precursor cell.

### Weighted region co-methylation network analysis (WRCMNA) reveals NDD microglial module

To further examine the impact of NDDs on regional changes in DNA methylation, we utilized a modified version of WGCNA based on similar approaches used to examine single CpG level methylation values from arrays (Horvath et al. 2012) to identify modules of co-methylation across the consensus background regions during the DMR analysis (Methods). After filtering for coverage and to remove low variance regions, 92,836 consensus regions were included in the network using all 49 WGBS samples (Figure S9) and 14 modules of co-methylation were identified (Table S12, Figure S10). The module eigengenes were analyzed for association with NDD diagnosis, sex, sequencing cohort, brain region, PMI and age (Figure 6A and Table S12). NDD diagnosis was significantly correlated with eigengenes from several modules; however, all but one module (purple) was also correlated with sequencing cohort, indicating that these modules were largely driven by the sequencing platform and may not be specific to NDD diagnosis. Several modules were also significantly correlated with sex and brain region (BA9 versus BA19).

**Figure 6.**
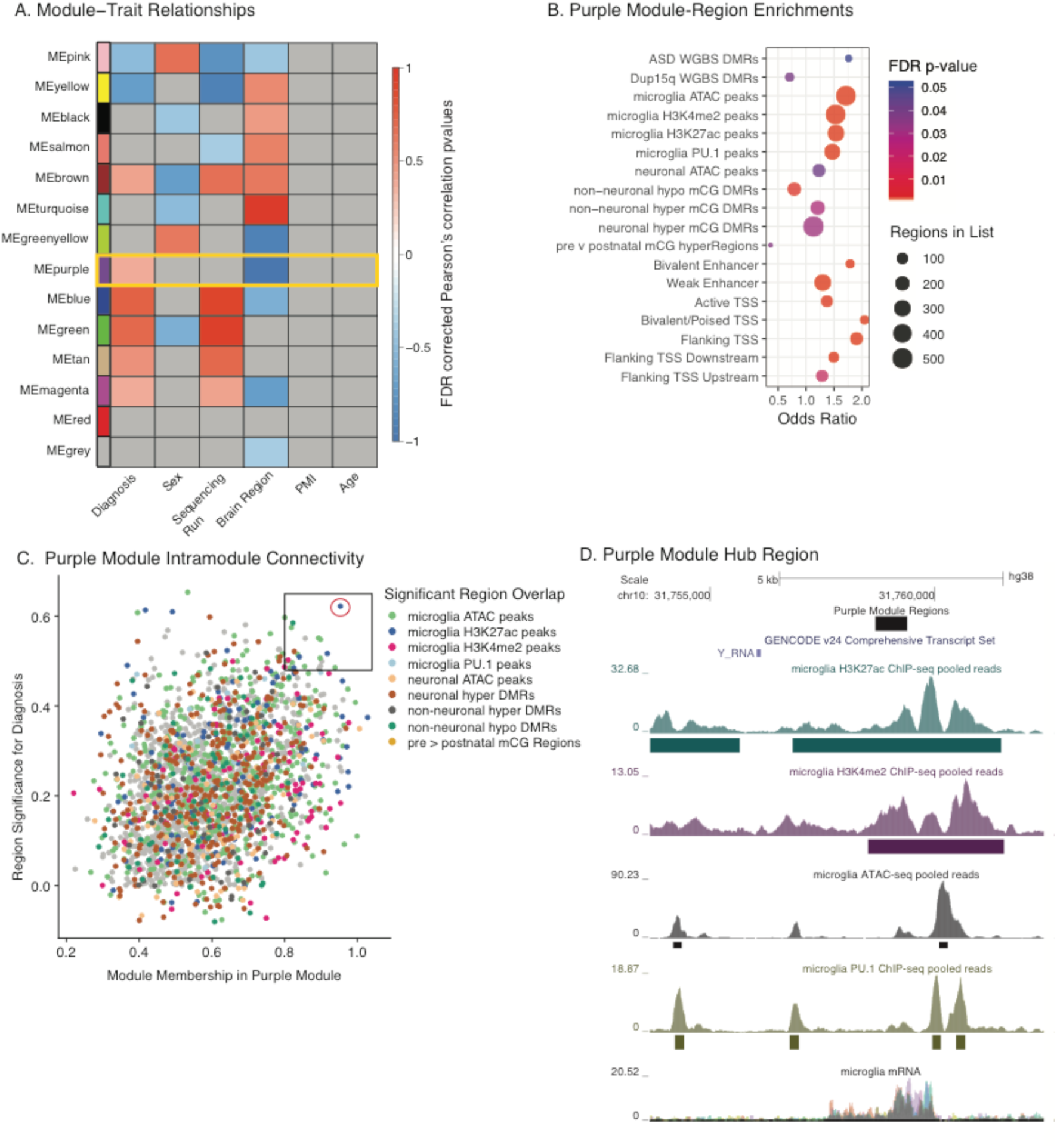
Weighted region co-methylation network analysis of NDD cortex reveals enrichment for a microglial regulatory module. **A.** The smoothed mCpG for each region identified in the consensus background regions from DMR analysis was used as input to a weighted and signed correlation network to identify 13 modules of co-methylation (and one non-correlated grey module) for all 49 WGBS samples. Module eigengenes shown in Figure S9 and Table S12. FDR corrected Pearson’s correlation p-values were used to define the relationship between module eigengenes (first principal component) and traits. Only the purple module (yellow box) was significant for diagnosis and brain regions without also being associated with sequencing run (FDR p<0.05). Diagnosis included variables for each NDD (ASD, Dup15q and RTT) and controls. Grey represents a nonsignificant correlation (FDR p>0.05). **B.** Regions within the purple module were enriched (Twotailed Fisher’s Exact Test) for numerous markers of microglial regulatory regions as well as NDD DMRs identified in Figure 1. ASD DMRs approached significance at FDR p< 0.051. All others FDR p<0.05. **C.** The Module Membership (absolute value of the correlation coefficient between smoothed methylation values and the Purple ME) is positively related to the Gene-Trait Significance values for Diagnosis (absolute value of the correlation coefficient for methylation and diagnosis status). The top 20% of intramodule connections are boxed and the top ranked hub region is circled in red. The regions that significantly overlap regulatory elements identified in B are color coded. Note the high prevalence of microglial regulatory regions in the hub gene region (box). **D**. The top hub region (circled in C) overlaps multiple microglial enhancer regions as shown in the UCSC genome browser. Microglial tracks are from http://cmm.ucsd.edu/qlass/qlasslab/quide.html (Gosselin et al. 2017). Statistics and modules are in Table S12.

The purple module that significantly correlated with NDD diagnosis and brain region but no other factor in the analysis, was further explored to identify potential impacts of altered DNA methylation within the regions contained in the module. Module regions (2346) were overlapped with regulatory regions identified from ENCODE chromHMM for PFC, regions of differential chromatin accessibility and methylation within NeuN+ neurons compared to NeuN-non-neurons (microglia, astrocytes, etc.) (Lister et al. 2013; Fullard et al. 2017), and regulatory regions identified from acutely isolated human brain microglia (Gosselin et al. 2017) (Table S12). The purple module was significantly (two-tailed Fisher’s exact test FDR p<0.05) enriched for markers of human microglial regulatory regions (PU.1 binding sites, open chromatin, and histone marks), neuronal open chromatin, enhancer and TSS markers, as well for as methylation markers of non-neuronal regulatory regions (Figure 6B). In addition, Dup15q DMRs identified in this study were significantly enriched and ASD DMRs approached significance for enrichment (FDR p<0.051) (Figure 6B and Table S12). The purple module was enriched for GO terms (Table S12) associated with cellular adhesion, cell-cell interactions, nervous system development, plasma membrane, channel activity and ion binding. Together, these enrichments indicate that regions within the purple module regulate a variety of cellular functions associated with cell-cell interactions during brain development.

We next identified top regulatory regions within the purple module by examining the level of intramodule connectivity (kME), a measure commonly used in WGCNA to identify hub genes that may serve as drivers of module expression (see methods). These regions of high intramodular node connectivity may serve as “hub” regions with key regulatory roles for the module. The top ranked hub regions for the purple module were significantly (two-tailed Fisher’s exact test FDR p<0.05) enriched for numerous markers of microglial regulatory activity including ATAC signal, histone marks for enhancer regions, and PU.1 binding sites (Figure 6C boxed region). For example, the top ranked region from the purple module (Figure 6C red circle) falls within an intergenic microglial regulatory region that is largely devoid of annotated genes, but enriched for human microglial expression and numerous enhancer marks, indicating it may be either a previously unannotated gene or an enhancer RNA expressed in microglia.

## Discussion

This unbiased epigenomic analysis of alterations in cortical DNA methylation across three distinct NDD disorders identified differentially methylated loci enriched for neuronal and glial regulatory regions associated with regulation of metabolic, transcriptional, and immune processes that were also highly enriched for genetic NDD risk variants. This is the first study to demonstrate overlap between epigenetic and genetic risk at a genome-wide level for NDDs, an observation that has been previously seen at candidate gene loci *MECP2* and *OXTR* (Nagarajan et al. 2008; Gregory et al. 2009) and the imprinted locus 15q11.2-q13.3 (Dunaway et al. 2016). The regions of differential methylation in NDDs indicate early developmental alterations in developmental gene expression across multiple cell types including neurons and microglia. Our findings leverage previous transcriptional and epigenomic profiling in human ASD brain that have consistently revealed both synaptic and immune dysfunction (Gupta et al. 2014; Nardone et al. 2014, 2017; Dunaway et al. 2016; Parikshak et al. 2016; Sun et al. 2016). ASD DMRs in particular were highly overlapped with microglial regulatory regions, and DMR associated genes from all three NDDs were enriched for genes regulated during microglial development, microglia from NDD mouse models, and other microglial inflammatory conditions. Weighted region co-methylation analysis also revealed that the module correlated with NDD diagnosis was enriched for microglial regulatory regions, including the top hub regions. Together with the immune gene expression signature observed across multiple ASD human brain cohorts (Gupta et al. 2014; Gandal et al. 2016; Parikshak et al. 2016), our data supports a convergent role for altered microglial function in NDDs. WGCNA across developmental time in human prefrontal cortex and mouse microglia indicates that epigenomic alterations in microglia may occur during mid to late prenatal development. This is also consistent with alterations in microglia morphology and density (Vargas et al. 2005; Morgan et al. 2010, 2012, 2014; Tetreault et al. 2012) previously observed in a subset of postmortem ASD brains. Maternal immune activation and dysregulation during pregnancy is also a risk factor for ASD (Atladottir, Thorsen, Ostergaard, et al. 2010; Atladottir, Thorsen, Schendel, et al. 2010; Zerbo et al. 2013; Lyall, Ashwood, et al. 2014; Lyall, Schmidt, et al. 2014). Children with ASD often display a variety of immune related abnormalities including altered cytokine expression (Goines and Ashwood 2013) and changes in immune cell populations (Ashwood et al. 2011), suggesting a shared disruption in the development of the prenatal immune system and brain.

As the resident immune cells in the brain, microglia may be a common cell type impacted by diverse NDD etiologies because they serve as important sentinels responding to both genetic and environmental disruptions. However, it remains unclear if alterations in DNA methylation in NDDs are a contributing cause or indirect consequence of the disorder and that the identified DMRs may serve as serve as a unique readout of both genetic and environmental perturbations to the developing brain. Microglia not only constantly monitor the brain for signs of infection, but also modulate synaptic transmission by secretion of neurotrophins (Parkhurst et al. 2013), regulate synapse formation and elimination (Tremblay et al. 2010; Paolicelli et al. 2011; Schafer et al. 2012; Zhan et al. 2014), and control neuronal precursor levels (Cunningham et al. 2013). Microglia also respond to genetic abnormalities that impair neuronal function. For example, multiple studies in mouse *Mecp2* mutant models of Rett syndrome show evidence of microglial activation and involvement in disease progression (Derecki et al. 2012; Cronk et al. 2015; Horiuchi et al. 2016). In addition, conditional deletion of CCCTC binding factor (CTCF) specifically in postnatal glutamatergic forebrain neurons in mice decreased neuronal dendritic spine density, and microglia in these animals adopt an abnormal morphology and up-regulate transcription of microglial inflammatory genes (Mcgill et al. 2017). Therefore, immune signatures observed across transcriptomic and epigenomic studies in human ASD brain may be partially driven by an immune response to abnormal neuronal processes that arise from genetic and epigenetic etiologies. Regardless of the initial cause of the NDD, microglia may be an appropriate target for therapy, since altered immune function may have profound impacts on neuronal development and ongoing brain function. Our analysis provides initial support for this premise as all three NDD DMR associated gene lists significantly overlapped genes normalized by minocycline treatment in adult mice that had received PolyI:C MIA in utero, suggesting that therapeutics that alter inflammation may be beneficial for resetting the microglial transcriptome and function in NDDs.

One limitation of this work is the relatively small sample size due to limited region-matched availability of human brain samples and still relatively high cost of WGBS. However, this work is similar or larger in sample size than previous DNA methylation microarray studies when considering the evaluation of a single brain region. This study also specifically examined DNA methylation differences that were independent of sex, allowing for the identification of epigenomic signatures relevant to both male and female ASD cases. Furthermore, we uniquely leveraged three distinct NDDs (ASD, RTT and Dup15q) to identify regions of co-methylation impacted by NDD diagnosis and regulated by markers of microglial regulatory regions. The average WGBS genome coverage is not sufficient to identify single CpG methylation differences, but does allow for the assignment of DMRs (Korthauer et al. 2018), which represent biologically relevant regional methylation differences (Vogel Ciernia and LaSalle 2016). The batch effects (Leek et al. 2010) observed between the different NDDs sequencing cohorts were largely mitigated by identifying DMRs within cohort and then comparing across NDDs and identifying WRCMNA modules independent of sequencing cohort. With the continued decrease in cost of sequencing technologies and increased brain bank advocacy, future work can more fully characterize additional NDDs brain samples by WGBS on a single sequencing platform with additional brain regions, disorders, and cell type specific sorting.

In conclusion, findings from this study reveal a critical epigenomic signature in NDD cortex that overlaps with known genetic and immune dysfunction in NDDs. Integration with multiple data sources identified both neuronal and microglial cell types and pathways as potential relevant therapeutic avenues that may be commonly dysregulated mediators at the interface of genetic and environmental risk factors.

## Declarations

### Ethics approval and consent to participate

Not applicable

### Consent for publication

Not applicable

### Availability of data and material

Previously published datasets for human ASD and Control BA9 as well as Dup15q and Control BA19 samples are available at GEO GSE81541. The human BrainSpan Data is available at: (http://www.brainspan.org/rnaseq/) (Miller et al. 2014). Microglial developmental RNAseq data is available at GEO GSE99622 (Hanamsagar et al. 2017). All analysis code is available on Github at https://github.com/kwdunaway, https://github.com/aciernia, https://github.com/ben-laufer/CpG_Me, https://github.com/ben-laufer/DM.R and https://qithub.com/hyeveon-hwanq.

## Footnotes

Author’s Contributions A.V.C. designed the research approach, performed bioinformatic and statistical analysis, and wrote the manuscript. B.I.L. developed the WGBS alignment and DMR calling pipeline, contributed to the bioinformatic analysis, and helped prepare the manuscript. K.W.D. prepared sequencing libraries and initial bioinformatics processing. H.H., C.E.M., and R.L.C provided additional bioinformatics analysis. D.H.Y assisted with experimental design and manuscript preparation. J.M.L. assisted with research design and manuscript preparation. All authors read, edited and approved of the final manuscript.

## Acknowledgements

The authors would like to thank Dr. Shreejoy Tripathy at the University of British Columbia for helpful discussion on the human single cell RNAseq data.

